# What do we know about the missing millions of Earth’s insect species and can we improve their collection: evidence from bark beetles?

**DOI:** 10.1101/2024.03.20.585838

**Authors:** Nigel E Stork, Michael J. W. Boyle, Carl Wardhaugh, Roger Beaver

**Author notes:** **Author Contributions:** N.E.S conceived the study and made the first draft of the manuscript, N.E.S and C.W. collected most of the samples and made the initial sorting to morphospecies, R.B. made final species assessments and compiled data on Australian species, M.B. carried out the statistical analysis. All authors contributed to the final manuscript. **Competing Interest Statement:** The authors have no competing interests.

## Abstract

Only 20% of the estimated five million species of insects on Earth are named despite over 240 years of taxonomy. Yet insects are poorly represented in protected area assessments, and insect declines are of concern globally. Here we explore how to increase the discovery of new species and understanding of this group through analysis of 10,097 tropical rainforest bark beetles (Scolytinae) from eight different ecological studies using beetles between 2000 and 2018 in the Australian Wet Tropics. Of the 107 species identified, 58 are undescribed: an increase of 35% on the 166 species known from Australia. As hypothesised, new species are significantly smaller, less abundant and less widespread than described species making them more extinction prone than named species. Rarefaction indicates doubling sampling would increase the number of species by 17. Flight Interception Traps (FIT) collected 84% of individuals and 98% of species confirming the effectiveness of a single sampling method for some beetles. Increased locations and collection from the canopy may sample further species rather than additional collecting methods. Scolytines are relatively well studied with a cadre of taxonomists at the forefront of using modern methods to resolve formerly intractable groups. These new species are more likely to be named than others in many other beetle groups where taxonomy has largely stalled. To increase species description rates and to avoid most species becoming extinct before being named, we call on taxonomists to use new character systems provided by DNA methods and to look at working with Artificial Intelligence tools.

**Significance Statement:** In an era of rapid biodiversity loss, current conservation decisions for insects will continue to be based on a small and almost certainly biased sample of the world’s biota until more species are named. We demonstrate how large-scale sampling can dramatically increase the number of species discovered for one group of beetles and how these undescribed species are significantly smaller, less abundant and less widespread than named species. The identification and determination of undescribed species is rarely possible except when taxonomic expertise is available, as in the present study. Addressing the insect taxonomic bottleneck and increasing the rate of description will require the adoption of new and developing tools.

## Introduction

The number of insect species on Earth has been estimated to be around 5 million species (1-4). With only 1 million species named and excluding those accidentally named more than once, it appears that about 80% are yet to be discovered (1). Since taxonomists have been naming species for over 240 years, questions remain as to why these have not been found and described. Where are these un-named species (5) and what is the likelihood of their discovery? Scheffers and colleagues suggested that many of the undiscovered species are small, difficult to find and have small geographic ranges. They also postulated that most missing species live in biodiversity hotspots. Here we provide light on these questions through the study of bark beetles (Coleoptera: Scolytinae) sampled over a 20-year period in Australian tropical rainforest.

Bark beetles in a broad sense comprise two subfamilies of the Curculionidae: Scolytinae and Platypodinae (6). The larger of these, Scolytinae has more than 6000 species described to date and the vast majority are tropical or subtropical (7). Some are true bark beetles breeding in the phloem of trees, but many species of ambrosia beetles tunnel into the wood. Almost all scolytines are associated with fungi, but ambrosia beetles have an obligatory symbiotic relationship with ambrosia fungi which are actively cultivated in the tunnels, and form the sole food of both adults and larvae (e.g. (8-11). Other scolytines breed in seeds and fruits, the pith of twigs and the stems of herbs (8). Bark beetles have received considerable attention in recent years due to their association with fungi, and their ability to invade non-native regions with a devastating impact on the survival and distribution of some temperate tree species, with serious ecological and economic consequences. For this reason, there has been perhaps more taxonomic effort on Scolytinae than many other groups of beetles.

We first ask what species of scolytine are currently known from Australia and from Queensland before assessing how well we have sampled this fauna. We hypothesise that species we discover to be new to science will on average be much smaller and less abundant than previously named species (5). We discuss whether these traits may make them more vulnerable to extinction than previously named species. We test whether using more collecting methods or more locations will reveal more new species. Finally, we make suggestions as to how best to improve our sampling and naming of the unknown species around the world as well as the implications of such efforts for insect conservation.

## Results

Our analysis indicates the known Australian scolytine bark beetle fauna is 166 species (Atlas of Living Australia accessed 6 September 2023, supplemented by unpublished data from RB). This includes species found only on Xmas I. and Norfolk I. and a few imported, but doubtfully established species. We have excluded incompletely determined species, and species not occurring in Australia such as those from New Zealand. Of the 166 species, 134, or 80.7% of the fauna are known to occur in Queensland. Eighteen species are introduced and 64 are endemic

A total of 10,097 bark beetles were collected in the eight studies and initially sorted to 80 morphospecies. After study by RB these were identified to 107 species of which 58 are considered to be undescribed species (Supplementary Table 1). Most new species are in tribes comprising mostly small and morphologically homogeneous species, such as Cryphalini, Trypophloeini and Ernoporini (until recently lumped in ‘Cryphalini’ (12) as well as Dryocoetini. Our list reflects current nomenclature (Supplementary Table 1).

FITs accounted for almost all of the individuals and species (Table 1, Figure 1) with only three species not being caught using this method: the Atherton Tableland fragmentation experiment contributing 2,935 individuals and 57 species, Atherton altitudinal transect 3,886 individuals and 46 species, and Paluma altitudinal transect contributing 469 individuals and 24 species. Daintree experiments in total contributed 2,806 individuals and 66 species with the ground FIT contributing the most in terms of 1156 individuals but only 27 species whereas the canopy FIT with 888 individuals produced 57 species.

**Table 1.** Summary of abundance, total number of species by a) location and b) sampling method (New = Percentage of collected species that are unnamed)

|  | Atherton<br>Tablelands | Atherton<br>Transect | Paluma<br>Transect | Daintree |
| --- | --- | --- | --- | --- |
| Individuals | 2934 | 3886 | 469 | 2805 |
| Species | 57 | 46 | 24 | 70 |
| Unique | 14 | 7 | 2 | 37 |
| Named | 41 | 32 | 19 | 39 |
| Unnamed | 16 | 14 | 5 | 31 |
| New | 28% | 30% | 21% | 44% |

b)
|  | Beating | FIT<br>Ground | FIT<br>Canopy | Fogging | Fruit | Leaf<br>Litter | Light |
| --- | --- | --- | --- | --- | --- | --- | --- |
| Individuals | 5 | 8507 | 985 | 1 | 231 | 139 | 226 |
| Species | 4 | 83 | 58 | 1 | 6 | 7 | 21 |
| Unique | 0 | 39 | 13 | 0 | 0 | 1 | 3 |
| Named | 3 | 50 | 34 | 1 | 4 | 5 | 17 |
| Unnamed | 1 | 33 | 24 | 0 | 2 | 2 | 4 |
| New | 25% | 40% | 41% | 0% | 33% | 29% | 19% |

**Figure 1.**
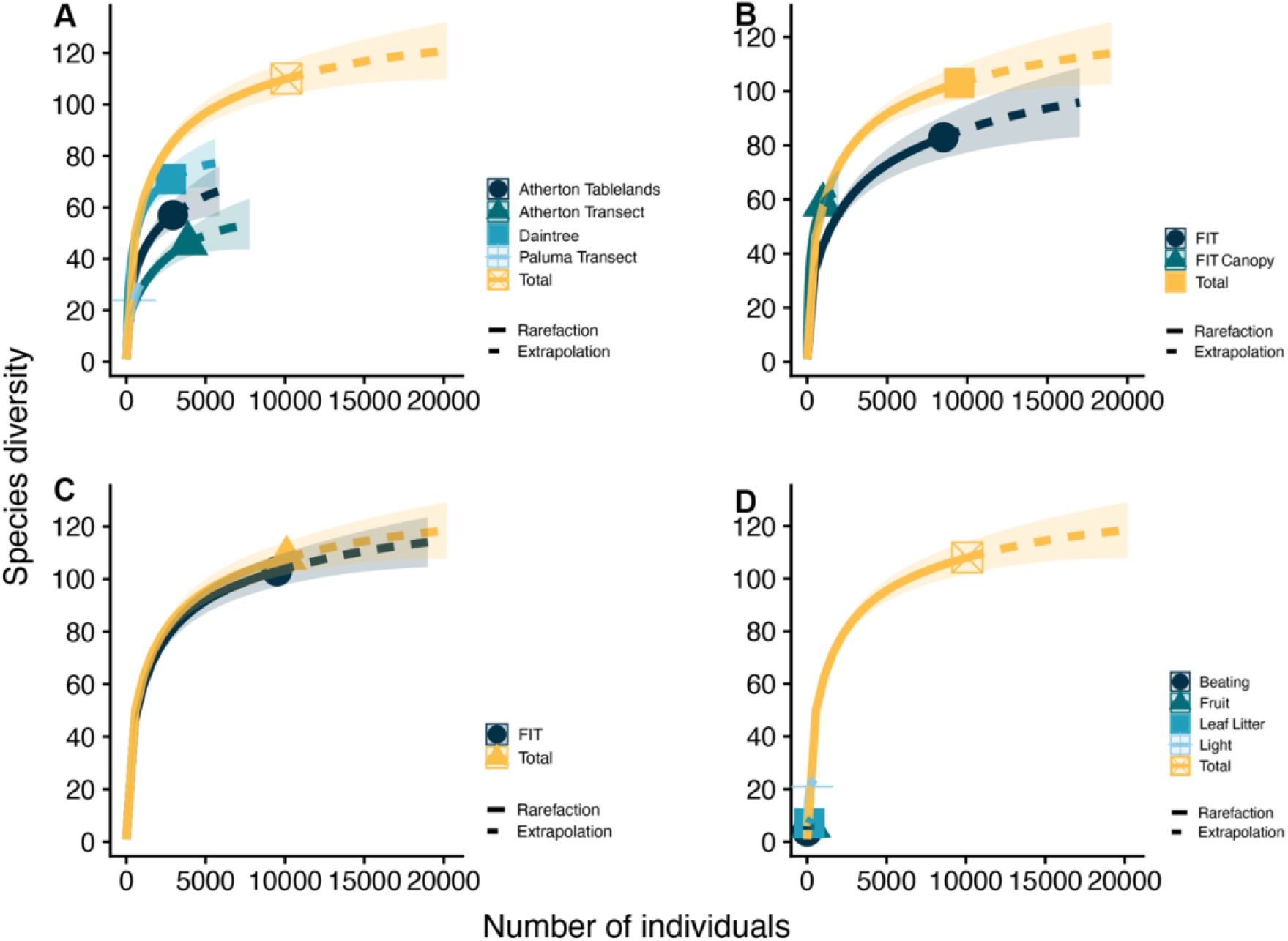
Sample based rarefaction of species richness estimates from **A)** sampling sites **B) FIT** sampling in ground and canopy layers C) Ground and canopy FIT combined and D) Other non-FIT sampling methods. In each case, the **total** estimation line represents the sum of either all sites or methods, showing that a combination of sites and strata increases species richness significantly.

Rarefaction indicated that our samples were representative of the community, with only 17 more species expected to be collected following a doubling of the samples (Fig. 2). Total observed species richness was 108, with an estimated richness of 124 (lower = 108, upper = 150). Daintree had the highest diversity as a proportion of the overall sampled community, followed by Atherton Transect, Atherton Tablelands and Paluma Transect (Fig. 1A). FIT traps on the ground contributed roughly 80% of the community, with the remaining 20% appearing only in canopy FIT traps (Fig. 1B). Together, samples from these FIT traps in both strata represented the entire extrapolated community (Fig. 1C), with other methods of collection contributing negligible numbers of species (Fig. 1D).

**Figure 2.**
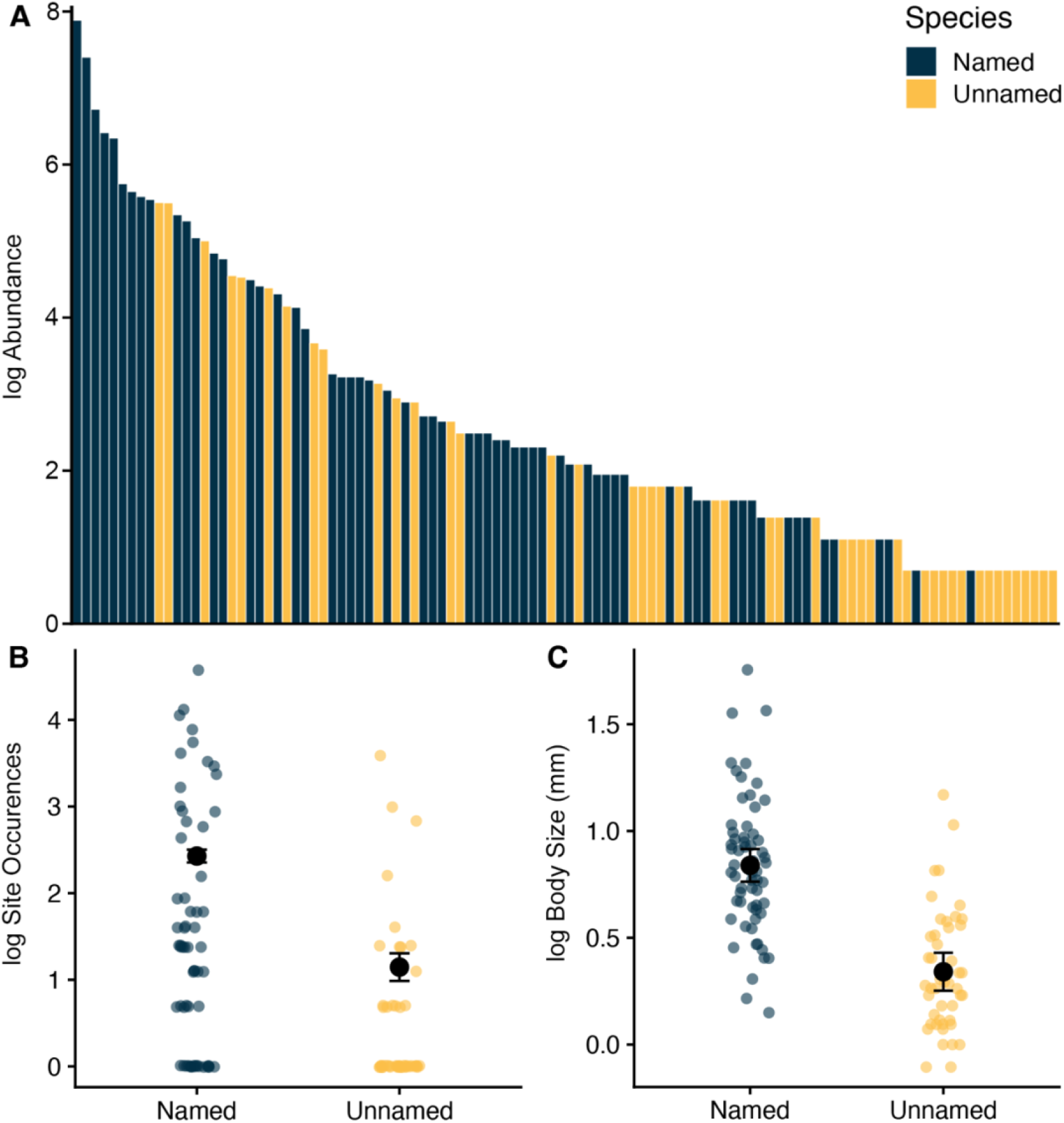
A) Log abundance of all bark beetle species from most to least abundant, B) Box and whiskers comparison of named and unnamed species across sites, and C) Box and whiskers comparison of named and unnamed species body sizes.

DCA analysis showed that our samples in general comprised one large meta-community of species, with few significant differences observed among communities sampled from each location (Supplementary Figure 1). While all levels in each DCA clustered clearly, only Daintree was significantly different from other sites in **DCAa** (Permanova: Atherton Tablelands vs Daintree *df* = 1, *F* = 5.23, *R*^*2*^ = 0.37, *p (adj*.) = 0.02. Daintree vs Paluma Transect *df* = 1, *F* = 6.19, *R*^*2*^ = 0.47, *p (adj*.) = 0.04) and no elevations were significantly distinct in **DCAb**. The canopy community was significantly distinct from the ground community in **DCAc** (Permanova: Canopy vs Ground *df* = 1, *F* = 2.24, *R*^*2*^ = 0.10, *p* = 0.02). The clustering of sites roughly corresponds with the clustering of elevations, with Daintree being the lowest elevation (50m), Paluma and Atherton Transects representing a range of elevation (100m – 1000m) and Atherton Tablelands representing a high elevation community (800m). The significant difference observed between Daintree and other sites may be due to the fact that Daintree includes canopy samples, which stood out significantly from ground level samples.

Named species were the most abundant species, and the singletons were almost all unnamed species, but in between there is some variation (Fig. 2a). The unnamed species are significantly smaller than the named species (ANOVA: *F*_1,106_ = 70.84, *p* < 0.0001, *R*^2^ = 0.40)(Fig. 2b), being 65% smaller (0.91 mm) on average. Unnamed species also occur at significantly fewer trap sites (GLM: *Z* = -14.29, *χ*^*2*^ = 259.8, *p* = <0.0001) (Fig. 2c), with named species expected to occur at 11 traps on average versus 3 for unnamed species.

## Discussion

Our study has increased the number of Scolytinae found in Australia by 58 or an additional 35% and of Queensland by 43%. More detailed taxonomic analysis including examination of type specimens for some previously named species, may reveal that a few are existing known species, but this still represents a remarkable increase in our understanding of the bark beetle fauna of Australia. Stork, *et al*. (13) previously found that 38.5% were undescribed species in a sample of 156 species across 21 families of beetles (14) suggesting our Scolytinae analysis may be typical for beetles as a whole in these samples. In this Discussion we focus on three main issues that arise from our analysis.

### Traits of undescribed species and their vulnerability to extinction

Species new to science are predicted to be on average smaller, less abundant and have narrower ranges than previously discovered species (5, 15). Our analysis confirmed this although some new species were surprisingly abundant (Fig. 1a,b) as often there can be a short-lived ‘pulse’ in numbers of individuals of a species with the mass emergence of bark and ambrosia beetles from a single dead tree or branch which lies within the attractive range of the trap.

Many undescribed species are likely to be local endemic species which may be less genetically able to adapt to a warmer or drier environment (16). It has also been predicted that most undescribed species are likely to be in the world’s 36 biodiversity hotspots (5) where less than 10% of natural intact vegetation remains (17). These hotspots are under pressure of further vegetation clearing due to an increasing demand for agricultural land (17, 18). It would seem that undescribed species more likely to become extinct than named (19) through the combined threats of land use change and climate change.

### How much sampling and what methods?

Our sampling was overwhelmingly by FIT and most species at all sites were collected by this effective method. Our rarefaction curves for all methods compared to FIT alone does indicate that using other methods generates more species but not by many more. Some methods such as rearing from fruit, though, provide valuable host-specificity information (20). Further, our analyses suggest that since smaller species tend to have narrower ranges additional locations would most likely produce further species rather than the use of other methods. The Wet Tropics, like other biodiversity hotspots around the world is home to many narrow range endemics usually on isolated mountain blocks (21). Similarly, the distinctiveness of canopy samples suggests more sampling in the canopy would be productive. We should not be surprised by the effectiveness of a single and simple sampling method such as FIT as others have shown. For example, Gauld collected 150 species of Pimpline Ichneumonid wasps in Costa Rica using standard Malaise traps. Local farmers emptied and maintained his traps at 17 sites throughout Costa Rica for a grand total of over 100 Malaise trap years between 1986 and 1990 (22). All additional sampling by hand collecting and by other collectors added no new species.

Elsewhere the introduction of novel sampling methods has surprised entomologists with their effectiveness at sampling previously poorly known faunas. Simple cloth Winkler bags extracting soil and leaf litter insects from sieved samples can be set up in field camps without light and heat sources and are highly effective at sampling soil insects such as Staphylinoidea and ants (23). Similarly, many taxonomists were excited to see canopy fogged knockdown insecticide samples for the first time in the early 1980s. Stork (24) and more than 30 specialist taxonomists sorted 24,000 arthropods from trees in Borneo to at least 3,000 species. Chalcidoidea comprised 739 species with only nine species having 10 or more individuals; almost certainly the vast majority were new to science (25). In Ecuadorian Amazonia rainforest fogging samples Scolytinae were well represented with an estimated 400 morphospecies from just 2,500 individuals (26). Stork and Grimbacher (14) used *IndVal* (27) to assess stratum specialisation and found 25% were canopy specialists, 25% were ground specialists, while the remaining 50% showed no stratum specialisation. For the Scolytinae seven species were canopy specialists and three were ground specialists. This shows that the canopy can have a quite different fauna to the ground level and sampling the canopy at the other sites may have produced further species.

FITs, Winkler bags and knockdown insecticides are examples of unbiased sampling methods which may accumulate numbers of individuals slowly but may give a representative selection of the species in an area. This is in contrast to biased sampling methods such as light traps and chemical lures which attract often large numbers of individuals of certain species. Having made that distinction, it should be noted that biased methods sometimes too can be useful for finding previously undiscovered species (28).

### Can we recognise, name and describe all new species?

For many groups of insects current morphological methods are unable to determine if a collection of individuals is one or many species and no number of additional years of study with the same approach will resolve them. Other character systems and tools including molecular tools are needed to resolve these taxonomically intractable groups and a collective approach is required with individuals who can bring together the tools needed to make taxonomic breakthroughs. *Hypothenemus eruditus* is typically identified as the world’s commonest bark beetle, but DNA analysis suggests it comprises many cryptic species and we are unable to associate morphological characters with true species limits (29). For Scolytinae there is a very active group of taxonomists who are using all available techniques to resolve uncertain groups and in doing so describe new species. The prospects are relatively good that the species we have discovered might be described therefore within the next 20 or so years. Furthermore, as with many insect groups, there are some scolytine taxa that seem to be taxonomically intractable, thus hindering effective taxonomic effort. The ‘Cryphalini’ were a particularly large, intractable group of bark beetles, but an extraordinary collective effort has shown that this comprises three tribes (Trypophloeini, Ernoporini, and Dryocoetini) and the taxonomic boundaries have been redefined (12) opening up the opportunities for further taxonomic revisions. It is no surprise then that 53 of the 58 new species we discovered are in these tribes. While we have focussed on collecting new specimens to increase discovery rates, we recognise that taxonomic revisions are frequently driven by the existence of large numbers of unresolved species, many collected many decades ago, but sitting in museums.

Scolytinae is just one of a number of subfamilies in the Curculionidae and there are 194 families of Coleoptera and close to 750 families in the five most species rich Orders of insects. The prospects of new species being described for many of these is much less because of the lack of expert taxonomists and because many groups within these are taxonomically intractable. Many families have had virtually no taxonomic effort on them for 100 years. These groups are unlikely to be known any better in the future without a change in the way taxonomy is done. Annual description rates for beetles and insects were roughly 1,050 and 7,200 species respectively, in the period 1978-1987 (30) and there is little evidence to suggest that this has changed much in recent decades.

In conclusion, we have shown that for bark beetles undescribed species are on average smaller, less abundant and occur at fewer locations and that intense sampling coupled with expert taxonomy can help improve their sampling. But what are the prospects for these ‘missing’ species? Eggleton (31) in his review of the state of the world’s insects suggests we need to resolve the insect taxonomic bottleneck, by finding ways of describing species faster and solving the intractable problems, such as incorporating DNA techniques into insect taxonomy using DNA barcoding and metabarcoding. Hulcr and colleagues (32) see a possible future where Artificial Intelligence machines have been trained for systematics and where robots become more efficient at identifying species than people. This may be achieved by taxonomists partnering with experts in AI and other modern tools. They urge that we must keep collecting to provide additional data and that taxonomists will still be vital to steer the decision-making processes.

We stress the economic importance of taxonomy as the essential basis for knowledge of pests, of invasive species, of integrated pest management, and taxonomists play vital roles in these fields of study. The planning for conservation of the world’s biota in protected areas is so frequently dependant on our knowledge of the distribution of plants and vertebrates. While we can assume that these ‘umbrella species’ will protect insects and other biodiversity we cannot be sure. Chowdhury, *et al*. (33) showed that 76% of 89,151 insect species assessed globally do not meet minimum target levels of protected area coverage and almost 2000 species do not overlap at all with protected areas. Species with low coverage occur in North America, Eastern Europe, South and Southeast Asia, and Australasia. We agree with their assessment that mapping important biodiversity areas must be upscaled to ensure nations capture insect diversity. Perhaps now is good time for taxonomy to step up, increasingly engage in modern technologies and qualify as big science (34, 35)?

## Materials and Methods

### Field sites and collecting methods

Approximately 30% of remaining rainforest in Australia is in a narrow 400km coastal strip and most is contained within the boundaries of the Wet Tropics of Queensland World Heritage Area (WHA). Our study sites were in both the lowlands and uplands (Fig. 3). The main lowland site was under or close to the Australian canopy crane (40 m asl, 16^0^17’S, 145^0^29’E), at the Daintree Rainforest Observatory in the Daintree region. The main upland location was the Atherton Tablelands (17°–17°30′S, 145°30′–145°45′E) a plateau about 700–850 m above sea level, 25-50 km from the coast, with (mostly) fertile, basalt-derived soils (36)). Across the landscape a strong spatial gradient in climate (particularly precipitation) creates complex vegetation patterns (37). While there are still extensive tracts of continuous forest, much of the upland rainforest on the Atherton Tablelands was converted to pasture 80–100 years ago, resulting in a landscape mosaic of rainforest remnants in a matrix that consists mainly of pasture (38, 39).

**Figure 3.**
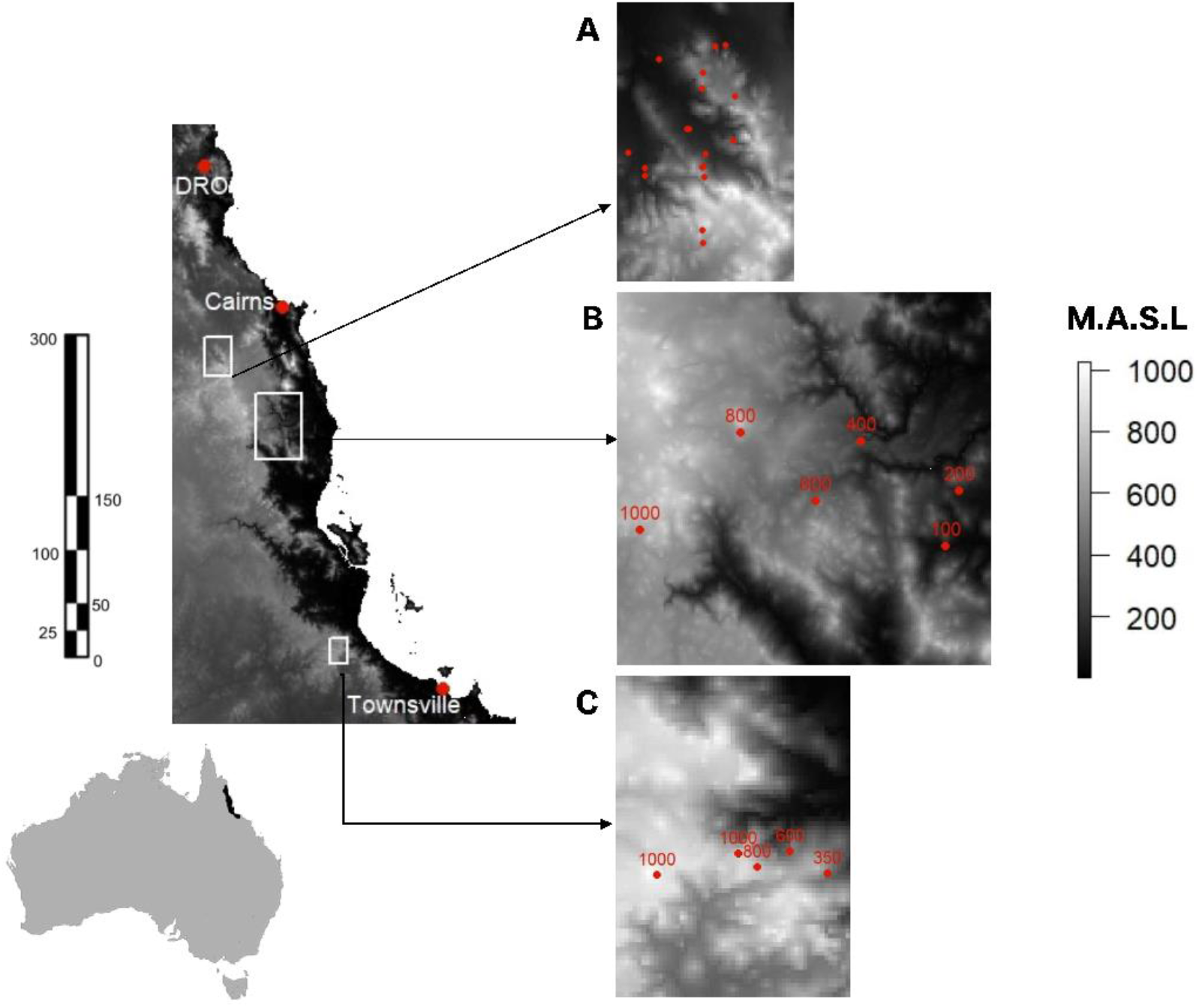
Map of the Wet Tropics of Queensland area showing inset boxes with field sites for three of the study areas (A - Atherton Tablelands fragmentation studies, B - Atherton elevational study, and C - Paluma elevational study) and a single point location for the fourth area, the Daintree Rainforest Observatory (DRO). Left-hand scale in kms, and righthand scale showing altitude in meters above sea level (M.A.S.L.). Based on map in (45). Figure created using packages *grid* (48) and *raster* (49) in R. Geoscience Australia data was used to develop mapping (50).

The eight studies where bark beetles were sampled are briefly described as follows (further details in Supplementary Information). Five studies were all carried out at the Daintree Rainforest Observatory with the other three located on the Atherton Tablelands (Fig. 3).

#### 1) Inter- and intra-seasonal variation

Insects were sampled using a Malaise trap modified to also act as a flight interception trap (FIT). Pairs of traps, one suspended 15–20 m above the ground in the canopy and one directly underneath on the ground, were located at five sites, each 40–60 m from each other. Traps were run for two weeks a month producing 45 monthly samples. (14)

#### 2) Distribution of beetles amongst canopy habitat types

Leaves, flowers, fruits and deadwood sampled for beetles in the upper canopy by beating with access provided by the canopy crane (40).

#### 3) Host specificity of beetles associated with fruit falls

Beetles were collected from fallen fruit of 18 species of tree once a month from October 2005 through to October 2010 (20).

#### 4) Vertical stratification of light-trapped beetles in the canopy

Beetles sampled by canopy light traps suspended 0 m, 10 m, 20 m and 30 m above the ground at five locations site in January and July 2012 (41).

#### 5) Temporal variation in leaf litter beetle abundance

The influence of climatic seasonality was quantified by sampling a fixed volume of litter monthly over 4 years and counting extracted beetles and ants (42).

#### 6&7) Atherton Tableland fragmentation studies

Bark beetles were sampled using small plastic FITs (see Supplementary Information) for 3 months from fragmented forests and larger intact blocks on the Atherton Tablelands (43). 3) The same plots were resampled six and 12 months after Cyclone Larry to examine the impact of cyclones on insects in fragmented forests (44).

#### 8) Elevational transects

Bark beetles were sampled with the same Malaise trap-based FITs used at the canopy crane site (45). Single traps were placed along two elevational transects on the eastern slopes, approximately 200 km apart, in the southern (Paluma transect at 350, 600, 800 m and two at 1,000 m asl) and central part Atherton transect at 100, 200, 400, 600, 800, and 1,000 m asl) of the Wet Tropics. Sampling was continuous from December 2006 to December 2007 with nine irregular length sampling periods.

### Beetle preparation and identification

All bark beetles were sorted to morphospecies, primarily by NES, CW and Peter Grimbacher. Most specimens, including all individuals of the most difficult groups, were identified by Roger Beaver, an expert on bark beetles. Species that were not named were considered to be new species based on his previous experience with Australian and S.E. Asian bark beetles. The specimens are housed at the Nathan Campus of Griffith University.

### Comparison with current Australian species lists

We used the current list of Australian species in the Atlas of Living Australia online resource as a baseline for our analysis of the names of species and their distribution. We particularly focussed on whether species had been recorded from Queensland with one of us (RB) adding to the current lists with his own data on species he had seen from Australia (R Beaver unpublished). We determined if species were likely to have been introduced or endemic.

### Statistical analysis

Sample based rarefaction was calculated using the package iNext (46) to visualise completeness of our samples and examine the projected species richness from different sites and collection methods. We also used Detrended Correspondence Analysis (DCA) to explore patterns of community assemblage using the package Vegan (47). Pairwise differences between sites were then compared using permutational analysis of variance (PERMANOVA) and p-values were adjusted using Hom-Bonferroni corrections to account for inflation of type-I error arising from multiple comparisons. For DCA and subsequent PERMANOVA only data from flight intercept traps were included.

ANOVA was used to test differences in the log mean body length of described and undescribed species, with five individuals of each species (or less if there were fewer available individuals) measured (to the nearest 0.1 mm) and then averaged as the mean body size for that species. We used a generalised linear model (GLM) to test for differences between named and unnamed species in the number of trap locations where each occurred, specifying a logistic error family (Poisson). P values for ANOVA were assessed using F statistics, and for GLM were assessed using Chi squared.

## Supporting information

Supporting information for Stork et al.

## Acknowledgements

We thank Jiri Hulcr and Ary Hoffmann for their comments on earlier drafts of this manuscript. N.E.S. is funded by the Australian Research Council grant DP200103100. We also thank Isaac Jennings and Queensland Cyber Infrastructure Foundation (QCIF) for assistance in the production of Fig. 1. We acknowledge the provision of data by Peter Grimbacher.

## Notes

### Competing Interest Statement

The authors have declared no competing interest.

