## Supporting information for Stork et al. for "What do we know about the missing millions of Earth’s insect species and can we improve their collection: evidence from bark beetles?"

Australia

^2^School of Biological Sciences, University of Hong Kong, Hong Kong

^3^Scion (New Zealand Forest Research Institute), 49 Sala Street, Rotorua, New Zealand

^4^161/2 Mu 5, Soi Wat Pranon, T.Donkaew, A.Maerim, Chiangmai 50180, Thailand

*Author for Correspondence: Nigel E Stork

**This PDF file includes:**

Supporting text: Details of the different sampling studies.

Table S1 List of all species sampled

Figure S1 Results of DCA

SI References

**Supporting text: Details of the different sampling studies.**

1) *Inter- and intra-seasonal variation*

Insects were sampled using a Townes-like Malaise trap modified to also act as a flight interception trap (FIT) with trays containing a 40% solution of propylene glycol placed either side of the base of the central net. Insects would either fly upwards and be caught in the Malaise bottle or fly/drop downwards and collect in the trays (1-3). Pairs of traps, one suspended 15–20 m in the canopy and one directly underneath on the ground, were located at five sites, each 40–60 m from each other. Traps were run for two weeks a month (with a few exceptions) from March 2000 to February 2004). This generated five replicate canopy FIT, canopy Malaise, ground FIT and ground Malaise samples, for each of the 45 monthly sampling periods.

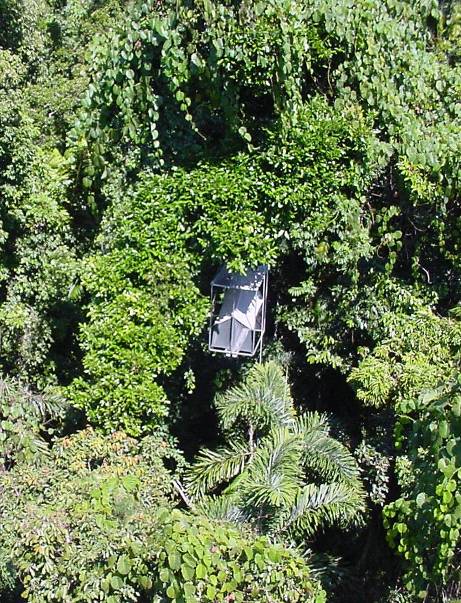

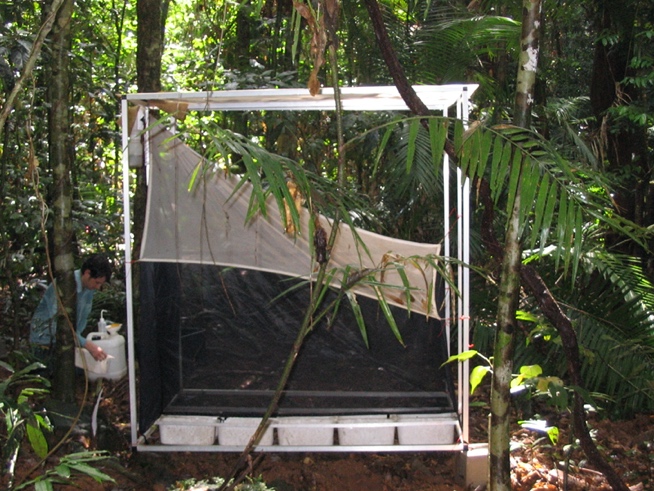

Malaise/FIT traps in the canopy and on the ground

*2) Distribution of beetles amongst canopy habitat types*

Using the canopy crane beetles were sampled monthly for 12 months by beating from five microhabitats; mature leaves, new leaves, flowers, fruit, and suspended dead wood from 23 locally common canopy plant species (4-8). One to three samples at a time depending on availability of microhabitat resource.

*3) Host specificity of beetles associated with fruit falls*

Fruit falls and beetles from fruits were sampled in situ and up to 10 min was spent at fruit

falls for a given plant species (9). During this time, fruits were placed over a white sheet and were shaken or broken apart to dislodge adult beetles that were subsequently collected using a pooter and stored in ethanol. Fruits at all available decomposition stages were sampled. Fruits were found and beetles were extracted on 55 different dates over a five-year period. There were some months where no fruit-associated beetles were sampled, although beetles were usually sampled from fruits of at least one plant species at each sampling date. Over 5 years, a total of 5157 individual beetles identified to 18 families and 73 species were collected in 94 samples from the fruits of 18 different plant species.

*4) Vertical stratification of light-trapped beetles in the canopy*

Light traps were suspended at different heights from ground level to 30 m in the canopy at four locations. Each night a trap was placed at a different height at each of the four locations and these were then rotated nightly for two weeks (10). Sampling was in January and July.

*5) Temporal variation in leaf litter beetle abundance*

Beetles within a standardised volume of litter were sampled each month over 4 years (January 2006–December 2009, except April 2006), resulting in 47 samples in total (11). Sampling involved a volume of five litres of sifted litter, collected from the ground. All litter was placed into a sifter (1 cm mesh size) to separate fine litter (and invertebrates) from coarse leaf and small woody material. Sampling continued until 5 litres of fine litter was collected. This material was then divided into equal subsamples and placed in a Tullgren funnel (internal diameter ~ 400 mm) for 12 h. All beetles were separated, counted and stored in ethanol.

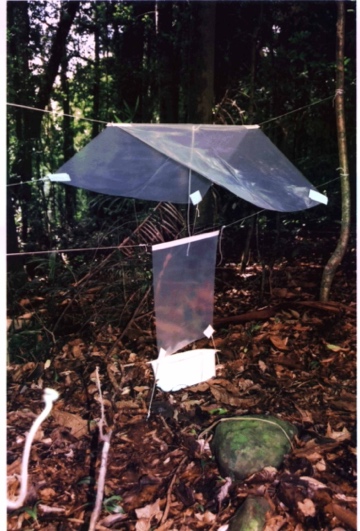
*6&7) Atherton Tableland fragmentation studies*

Original study:

Small ‘window trap’ like FITs. Six replicate sites in each of four habitat categories: pasture, small rain-forest remnants, edges and centre of large rainforest remnants (12-15). Four FITs at each site and sampled for 14 days. Note these FIT are about a third of the size of those used as part of a Malaise trap.

Second study following cyclone Larry (16):

Six replicate sites in each of three habitat categories: small rain-forest remnants, edges and centres of large rainforest remnants. Four FITs at each site and sampled for 14 days twice in the year.

*8) Elevational transects*

*Atherton altitudinal transect*

One Malaise trap used as an FIT as in the Canopy crane inter- and intra-seasonal study

at each of six elevations (100, 200, 400, 600, 800, 1000m) (17). Running continuously through December 2006 to December 2007 with samples collected at eight irregular intervals from. Access prevented a few samples not being collected for several of the sites. So roughly an average of 300 days sampling per trap:

*Paluma altitudinal transect*

One Malaise trap used as an FIT as in the Canopy crane inter- and intra-seasonal study

at each of six elevations (350, 600, 800, 1000m (two sites at 1000m)) collecting only as an FIT (17). Running continuously through December 2006 to December 2007 with samples collected at eight irregular intervals from. Access prevented a few samples not being collected for several of the sites. So roughly an average of 300 days sampling per trap:

**Supplementary Table 1 List of all species sampled, mean body size and abundance according to sampling method (blue) and location (yellow). Status: N named, U unnamed; Beat – beating, FIT Grnd – FIT Ground, FIT Canpy – FIT Canopy, Leaf – Leaf litter, Light – light trapping; Ather Tlands – Atherton Tablelands, Ather Trans – Atherton Transect, Pal Trans – Paluma Transect, Dain – Daintree.**

|  |  |  | Size (mm) | Beat | FIT Grnd | FIT Canpy | Fogg | Fruit | Leaf | Light | Ather Tlands | Ather Trans | Pal Trans | Daint |
| --- | --- | --- | --- | --- | --- | --- | --- | --- | --- | --- | --- | --- | --- | --- |
|  | Species | Status |  |  |  |  |  |  |  |  |  |  |  |  |
| Cryphalini | Cryphalus sp. 1 | U | 1.1 | 0 | 1 | 0 | 0 | 0 | 0 | 0 | 0 | 1 | 0 | 0 |
|  | Cryphalus sp. 2 | U | 1.075 | 0 | 0 | 4 | 0 | 0 | 0 | 0 | 0 | 0 | 0 | 4 |
|  | Cryphalus sp. 3 | U | 2.26 | 0 | 9 | 2 | 0 | 0 | 0 | 0 | 9 | 0 | 0 | 2 |
|  | Cryphalus sp. 4 | U | 1.2 | 0 | 0 | 2 | 0 | 0 | 0 | 0 | 0 | 0 | 0 | 2 |
|  | Cryphalus sp. 5 aff. brimblecombei (Schedl) | N | 1.6 | 0 | 0 | 61 | 0 | 0 | 0 | 0 | 0 | 0 | 0 | 61 |
|  | Cryphalus sp. 6 aff. ovalicollis (Schedl) | N | 1.8 | 0 | 825 | 0 | 0 | 0 | 0 | 0 | 722 | 103 | 0 | 0 |
|  | Cryphalus sp. 7 | U | 1.32 | 0 | 241 | 2 | 0 | 0 | 0 | 0 | 239 | 2 | 0 | 2 |
|  | Cryphalus sp. 8 | U | 1.78 | 1 | 0 | 21 | 0 | 0 | 0 | 0 | 0 | 0 | 0 | 22 |
|  | Cryphalus sp. 9 | U | 1.2 | 0 | 5 | 0 | 0 | 0 | 0 | 0 | 0 | 2 | 3 | 0 |
|  | Cryphalus sp. 10 | U | 1.3 | 0 | 1 | 1 | 0 | 0 | 0 | 0 | 0 | 1 | 0 | 1 |
|  | Cryphalus sp. 11 | U | 1.48 | 0 | 56 | 0 | 0 | 0 | 0 | 6 | 52 | 2 | 2 | 6 |
|  | Cryphalus sp. 14 | U | 1.26 | 0 | 3 | 5 | 0 | 0 | 0 | 0 | 1 | 1 | 0 | 6 |
|  | Cryphalus sp. 15 | U | 1.4 | 0 | 5 | 0 | 0 | 0 | 0 | 0 | 1 | 4 | 0 | 0 |
|  | Cryphalus sp. 16 | U | 1.3 | 0 | 0 | 2 | 0 | 0 | 0 | 0 | 0 | 0 | 0 | 2 |
|  | Cryphalus sp. 17 | U | 1.15 | 0 | 1 | 0 | 0 | 0 | 0 | 1 | 1 | 0 | 0 | 1 |
|  | Cryphalus sp. 18 | U | 1.12 | 0 | 5 | 0 | 0 | 0 | 0 | 0 | 4 | 1 | 0 | 0 |
|  | Cryphalus sp. 19 | U | 0.9 | 0 | 1 | 0 | 0 | 0 | 0 | 0 | 0 | 1 | 0 | 0 |
|  | Cryphalus sp. 20 | U | 1 | 0 | 1 | 0 | 0 | 0 | 0 | 0 | 0 | 0 | 0 | 1 |
|  | Cryphalus sp. 21 | U | 1.66 | 0 | 4 | 34 | 0 | 0 | 0 | 0 | 0 | 0 | 0 | 38 |
| Dryocoetini | Coccotrypes carpophagus (Hornung) | N | 1.92 | 0 | 192 | 34 | 0 | 4 | 7 | 74 | 103 | 5 | 15 | 188 |
|  | Coccotrypes cyperi (Beeson) | N | 2.24 | 0 | 317 | 101 | 0 | 87 | 61 | 0 | 29 | 125 | 13 | 399 |
|  | Coccotrypes queenslandi (Schedl) | N | 2.06 | 0 | 0 | 0 | 0 | 9 | 0 | 1 | 0 | 0 | 0 | 10 |
|  | Coccotrypes sp. AU2 nec advena Blandford | U | 1.8 | 0 | 1 | 1 | 0 | 55 | 36 | 0 | 0 | 0 | 0 | 93 |
|  | Coccotrypes vulgaris aff. (Eggers) | N | 2.14 | 0 | 9 | 0 | 0 | 0 | 0 | 0 | 9 | 0 | 0 | 0 |
|  | Cyrtogenius sp. 1 | U | 2 | 0 | 0 | 0 | 0 | 0 | 0 | 1 | 0 | 0 | 0 | 1 |
|  | Cyrtogenius sp. 2 | U | 1 | 0 | 1 | 0 | 0 | 0 | 0 | 0 | 0 | 1 | 0 | 0 |
|  | Cyrtogenius sp. 3 | U | 1.73 | 0 | 0 | 5 | 0 | 0 | 0 | 0 | 0 | 0 | 0 | 5 |
|  | Dryocoetiops moestus (Blandford) | N | 2.52 | 0 | 3 | 14 | 0 | 0 | 0 | 0 | 0 | 0 | 3 | 14 |
| Ernoporini | Eidophelus ater? (Schedl) | N | 1.5 | 0 | 1 | 0 | 0 | 0 | 0 | 0 | 1 | 0 | 0 | 0 |
|  | Eidophelus darwini (Eichhoff) | N | 2.64 | 0 | 1 | 4 | 0 | 0 | 0 | 0 | 0 | 1 | 0 | 4 |
|  | Eidophelus darwini aff. (Eichhoff) | N | 1.95 | 0 | 2 | 0 | 0 | 0 | 0 | 0 | 1 | 0 | 1 | 0 |
|  | Eidophelus sp. 1 | U | 1.36 | 0 | 5 | 5 | 0 | 0 | 0 | 3 | 2 | 3 | 0 | 8 |
|  | Eidophelus sp. 2 | U | 1.6 | 0 | 17 | 0 | 0 | 0 | 0 | 0 | 6 | 11 | 0 | 0 |
|  | Eidophelus sp. 3 | U | 1.34 | 0 | 147 | 0 | 0 | 0 | 0 | 0 | 147 | 0 | 0 | 0 |
|  | Eidophelus sp. 4 | U | 1.5 | 0 | 1 | 0 | 0 | 0 | 0 | 0 | 0 | 0 | 1 | 0 |
|  | Eidophelus sp. 5 | U | 1.1 | 0 | 1 | 2 | 0 | 0 | 0 | 0 | 0 | 0 | 0 | 3 |
|  | Eidophelus sp. 6 | U | 1.1 | 0 | 1 | 0 | 0 | 0 | 0 | 0 | 0 | 1 | 0 | 0 |
|  | Eidophelus sp. 7 | U | 1.8 | 0 | 1 | 0 | 0 | 0 | 0 | 0 | 0 | 1 | 0 | 0 |
|  | Eidophelus sp. 8 | U | 1.1 | 0 | 1 | 0 | 0 | 0 | 0 | 0 | 0 | 0 | 0 | 1 |
|  | Eidophelus sp. 9 | U | 1.82 | 0 | 0 | 4 | 0 | 75 | 0 | 0 | 0 | 0 | 0 | 79 |
|  | Eidophelus sp. 10 | U | 1.26 | 0 | 0 | 5 | 0 | 0 | 0 | 0 | 0 | 0 | 0 | 5 |
|  | Eidophelus sp. 11 | U | 1.075 | 0 | 0 | 4 | 0 | 0 | 0 | 0 | 0 | 0 | 0 | 4 |
|  | Eidophelus sp. 12 | U | 1 | 0 | 1 | 0 | 0 | 0 | 0 | 0 | 0 | 0 | 1 | 0 |
|  | Eidophelus sp. 13 | U | 1.5 | 0 | 0 | 1 | 0 | 0 | 0 | 0 | 0 | 0 | 0 | 1 |
|  | Eidophelus sp. 14 | U | 0.9 | 0 | 2 | 0 | 0 | 0 | 0 | 0 | 2 | 0 | 0 | 0 |
|  | Eidophelus sp. 15 | U | 1.33 | 0 | 3 | 0 | 0 | 0 | 0 | 0 | 3 | 0 | 0 | 0 |
|  | Eidophelus sp. 16 | U | 1.12 | 0 | 242 | 0 | 0 | 0 | 0 | 0 | 242 | 0 | 0 | 0 |
|  | Eidophelus tricolor (Lea) | N | 1.24 | 0 | 5 | 2 | 0 | 0 | 0 | 0 | 5 | 0 | 0 | 2 |
| Hylesinini | Ficicis despectus (Walker) | N | 2.4 | 0 | 13 | 10 | 0 | 0 | 0 | 1 | 4 | 3 | 1 | 16 |
|  | Ficicis elongatus (Schedl) | N | 1.9 | 0 | 3 | 0 | 0 | 0 | 0 | 0 | 3 | 0 | 0 | 0 |
|  | Ficicis maculipennis (Schedl) | N | 2.08 | 2 | 0 | 0 | 0 | 0 | 0 | 6 | 0 | 0 | 0 | 8 |
|  | Ficicis varians Lea | N | 2.04 | 0 | 25 | 0 | 0 | 0 | 0 | 0 | 24 | 0 | 1 | 0 |
| Hylurgini | Chaetoptelius tricolor (Schedl) | N | 2.54 | 0 | 20 | 0 | 0 | 0 | 0 | 0 | 20 | 0 | 0 | 0 |
| Hyorrhynchini | Sueus niisimai (Eggers) | N | 1.85 | 0 | 2 | 0 | 0 | 0 | 0 | 0 | 0 | 1 | 1 | 0 |
| Phloesinini | Hyledius cribratus (Blandford) | N | 2.58 | 0 | 18 | 5 | 0 | 0 | 0 | 0 | 1 | 17 | 0 | 5 |
| Trypophloeini | Hypothenemus areccae (Hornung) | N | 1.36 | 1 | 0 | 23 | 0 | 0 | 0 | 0 | 0 | 0 | 0 | 24 |
|  | Hypothenemus birmanus (Eichhoff) | N | 2.25 | 0 | 0 | 5 | 0 | 0 | 0 | 0 | 0 | 0 | 0 | 5 |
|  | Hypothenemus birmanus aff. (Eichhoff) | U | 1.92 | 0 | 5 | 13 | 0 | 0 | 0 | 0 | 3 | 0 | 0 | 15 |
|  | Hypothenemus crudiae (Eichhoff) | N | 1.575 | 0 | 4 | 0 | 0 | 0 | 0 | 0 | 4 | 0 | 0 | 0 |
|  | Hypothenemus eruditus cx Westwood | N | 1.16 | 0 | 0 | 5 | 0 | 0 | 1 | 0 | 0 | 0 | 0 | 6 |
|  | Hypothenemus seriatus cx (Eichhoff) | N | 1.56 | 0 | 116 | 0 | 0 | 0 | 0 | 0 | 116 | 0 | 0 | 0 |
|  | Hypothenemus sp. 3 | U | 1.26 | 0 | 5 | 2 | 0 | 0 | 0 | 0 | 4 | 0 | 0 | 3 |
|  | Hypothenemus sp. 7 | U | 1.3 | 0 | 0 | 0 | 0 | 0 | 1 | 0 | 0 | 0 | 0 | 1 |
|  | Hypothenemus sp. 8 | U | 1.4 | 0 | 0 | 1 | 0 | 0 | 0 | 0 | 0 | 0 | 0 | 1 |
| Xyleborini | Amasa fulgens (Schedl)(sp.2) | N | 2.8 | 0 | 0 | 2 | 0 | 0 | 0 | 0 | 0 | 0 | 0 | 2 |
|  | Amasa schlichii (Stebbing) | N | 3.6 | 0 | 1 | 0 | 0 | 0 | 0 | 0 | 1 | 0 | 0 | 0 |
|  | Amasa sp. 1 aff. truncata (Erichson) | N | 2.55 | 0 | 4 | 0 | 0 | 0 | 0 | 0 | 1 | 1 | 2 | 0 |
|  | Amasa sp. 3 aff. exacta (Schedl) | N | 2.16 | 0 | 1 | 8 | 0 | 0 | 0 | 2 | 0 | 1 | 0 | 10 |
|  | Amasa truncata (Erichson) | N | 3.4 | 0 | 46 | 0 | 0 | 0 | 0 | 0 | 29 | 8 | 9 | 0 |
|  | Amasa sp 4 aff. truncata (Erichson) | N | 3.175 | 0 | 4 | 0 | 0 | 0 | 0 | 0 | 4 | 0 | 0 | 0 |
|  | Ambrosiodmus asperatus (Blandford) | N | 2.46 | 0 | 5 | 6 | 0 | 0 | 0 | 0 | 1 | 2 | 1 | 7 |
|  | Ambrosiophilus australis (Schedl) | N | 2.2 | 0 | 4 | 0 | 0 | 0 | 0 | 0 | 2 | 2 | 0 | 0 |
|  | Ambrosiophilus celsoides (Hagedorn) | N | 3.73 | 0 | 3 | 0 | 0 | 0 | 0 | 0 | 3 | 0 | 0 | 0 |
|  | Ambrosiophilus compressus (Lea) | N | 3.22 | 0 | 187 | 4 | 0 | 0 | 0 | 0 | 69 | 100 | 5 | 17 |
|  | Ambrosiophilus latecompressus (Schedl) | N | 3.74 | 0 | 4 | 0 | 0 | 0 | 0 | 0 | 0 | 4 | 0 | 0 |
|  | Ambrosiophilus pityogenes (Schedl) | N | 2.6 | 0 | 2 | 1 | 0 | 0 | 0 | 3 | 1 | 1 | 0 | 4 |
|  | Ambrosiophilus sp. n.? | U | 1.75 | 0 | 0 | 1 | 0 | 0 | 0 | 0 | 0 | 0 | 0 | 1 |
|  | Beaverium annexus (Schedl) | N | 4.78 | 0 | 1 | 5 | 0 | 0 | 0 | 0 | 0 | 0 | 0 | 6 |
|  | Beaverium insulindicus (Eggers) | N | 5.78 | 0 | 8 | 1 | 0 | 0 | 0 | 0 | 1 | 3 | 0 | 5 |
|  | Cryptoxyleborus subnaevus | N | 2.5 | 0 | 0 | 8 | 0 | 0 | 0 | 1 | 0 | 0 | 0 | 9 |
|  | Debus pumilis (Eggers) | N | 2.08 | 0 | 197 | 7 | 0 | 0 | 0 | 3 | 8 | 187 | 0 | 12 |
|  | Diuncus haberkorni (Eggers) | N | 2.16 | 0 | 73 | 0 | 0 | 0 | 0 | 0 | 2 | 48 | 23 | 0 |
|  | Diuncus quadrispinosulus (Eggers) | N | 1.6 | 0 | 0 | 0 | 0 | 0 | 0 | 2 | 0 | 0 | 0 | 2 |
|  | Eccoptopterus spinosus (Olivier) | N | 2.7 | 0 | 4 | 5 | 0 | 0 | 0 | 0 | 2 | 2 | 0 | 5 |
|  | Eggersanthus sublaevis (Eggers) | N | 2.26 | 0 | 3 | 88 | 0 | 0 | 0 | 0 | 0 | 0 | 0 | 91 |
|  | Euwallacea destruens (Blandford) | N | 4.72 | 0 | 109 | 12 | 0 | 0 | 0 | 4 | 3 | 48 | 1 | 73 |
|  | Euwallacea fornicatus (Eichhoff) | N | 2.78 | 0 | 14 | 0 | 0 | 0 | 0 | 0 | 11 | 3 | 0 | 0 |
|  | Euwallacea funereus (Lea) | N | 3.5 | 0 | 0 | 3 | 0 | 0 | 0 | 0 | 0 | 0 | 0 | 3 |
|  | Euwallacea piceus (Motschulsky) | N | 2.24 | 0 | 250 | 13 | 0 | 0 | 0 | 0 | 245 | 0 | 0 | 18 |
|  | Euwallacea similis (Ferrari) | N | 3.04 | 0 | 2355 | 278 | 1 | 1 | 0 | 5 | 560 | 1295 | 312 | 473 |
|  | Fraudatrix pileatula (Schedl) | N | 2.68 | 0 | 0 | 10 | 0 | 0 | 0 | 0 | 0 | 0 | 0 | 10 |
|  | Gen. n. sp. n. | U | 3.22 | 0 | 2 | 33 | 0 | 0 | 0 | 0 | 0 | 0 | 0 | 35 |
|  | Microperus diversicolor (Eggers) | N | 1.96 | 0 | 2 | 22 | 0 | 0 | 0 | 0 | 0 | 0 | 0 | 24 |
|  | Microperus eucalypticus (Schedl) | N | 1.88 | 0 | 281 | 0 | 0 | 0 | 0 | 0 | 32 | 217 | 32 | 0 |
|  | Microperus sp. 1 aff. fulvulus (Schedl) | U | 2.8 | 0 | 1 | 0 | 0 | 0 | 0 | 0 | 0 | 0 | 0 | 1 |
|  | Microperus sp. 2 aff. quercicola (Eggers) | U | 1.67 | 0 | 2 | 1 | 0 | 0 | 0 | 0 | 1 | 0 | 1 | 1 |
|  | Planiculus bicolor (Blandford) | N | 2.34 | 0 | 11 | 0 | 0 | 0 | 0 | 77 | 0 | 11 | 0 | 77 |
|  | Xyleborinus artestriatus (Eichhoff) | N | 2.48 | 0 | 0 | 1 | 0 | 0 | 0 | 5 | 0 | 0 | 0 | 6 |
|  | Xyleborinus exiguus (Walker) | N | 1.74 | 0 | 5 | 6 | 0 | 0 | 0 | 2 | 3 | 2 | 0 | 8 |
|  | Xyleborus affinis Eichhoff | N | 2.28 | 0 | 235 | 0 | 0 | 0 | 1 | 17 | 92 | 143 | 0 | 18 |
|  | Xyleborus bispinatus Eichhoff | N | 2.62 | 0 | 9 | 1 | 0 | 0 | 0 | 4 | 4 | 4 | 0 | 6 |
|  | Xyleborus perforans (Wollaston) | N | 2.32 | 0 | 1520 | 69 | 0 | 0 | 32 | 8 | 53 | 1173 | 9 | 394 |
|  | Xylosandrus abruptulus (Schedl) | N | 1.94 | 0 | 153 | 0 | 0 | 0 | 0 | 0 | 2 | 128 | 23 | 0 |
|  | Xylosandrus monteithi Dole & Beaver | N | 3.14 | 0 | 81 | 0 | 0 | 0 | 0 | 0 | 6 | 75 | 0 | 0 |
|  | Xylosandrus morigerus (Blandford) | N | 1.8 | 1 | 597 | 9 | 0 | 0 | 0 | 0 | 38 | 139 | 7 | 423 |
|  | Xylosandrus queenslandi Dole & Beaver | N | 1.72 | 0 | 3 | 0 | 0 | 0 | 0 | 0 | 1 | 2 | 0 | 0 |
| Xyloctonini | Scolytomimus philippinensis Eggers | N | 2.36 | 0 | 2 | 5 | 0 | 0 | 0 | 0 | 0 | 0 | 0 | 7 |
|  | Scolytomimus pusillus (Eggers) | N | 1.5 | 0 | 5 | 6 | 0 | 0 | 0 | 0 | 1 | 0 | 2 | 8 |

**Supplementary Figure 1**

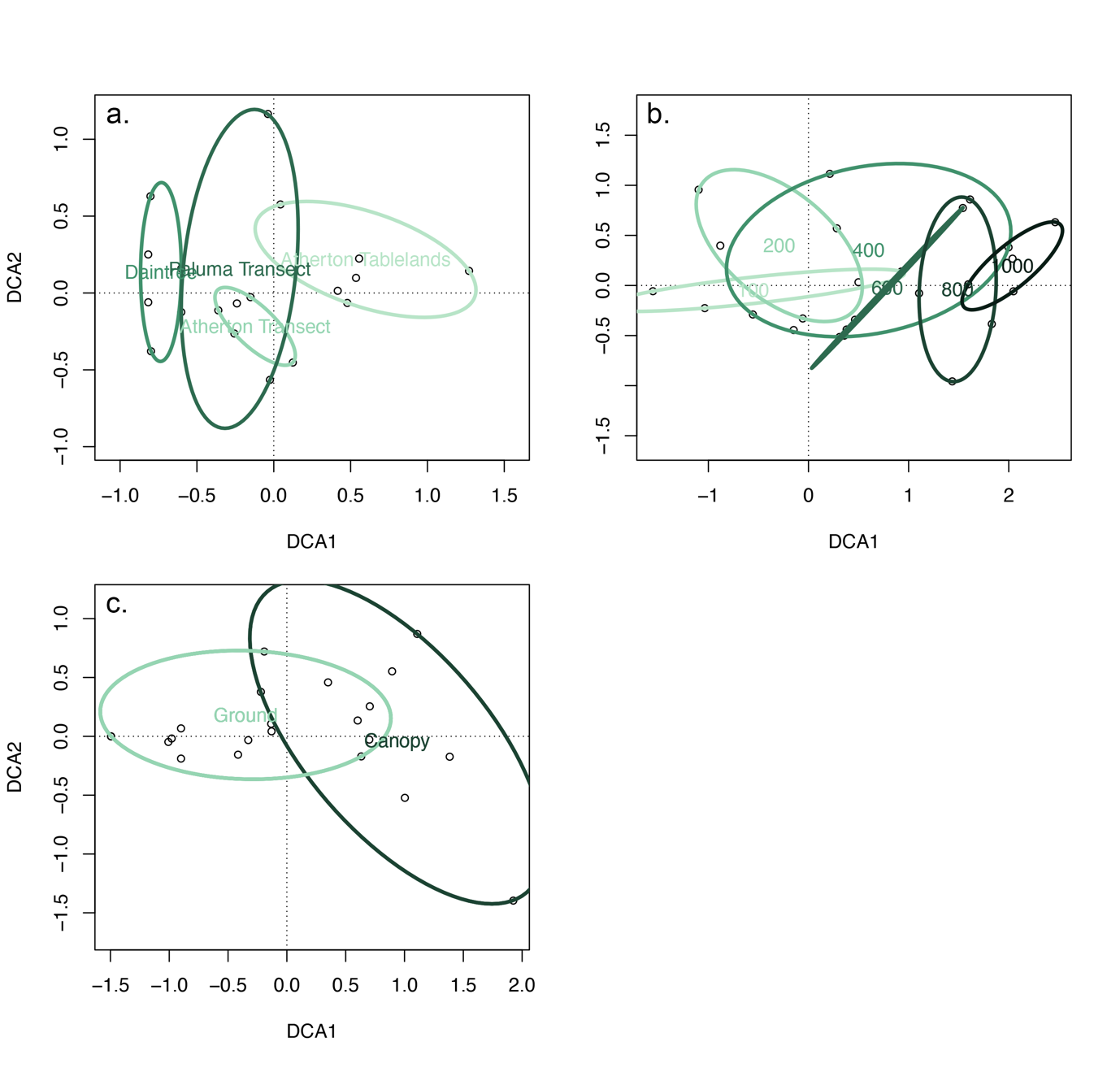

**Fig. 1 –** Detrended correspondence analysis (DCA) showing a visualisation of community level differences among **a.** Sampling sites, **b.** Sampling elevations (Atherton Transect only), and **c.** Sampling strata (Daintree only).
